# Distinct metabolic profiles of circulating plasmacytoid dendritic cells in systemic sclerosis patients stratified by clinical phenotypes

**DOI:** 10.1101/2024.11.27.625448

**Authors:** Beatriz H. Ferreira, Carolina Mazeda, Eduardo Dourado, João L. Simões, Ana Rita Prata, Rafael J. Argüello, Iola F. Duarte, Philippe Pierre, Catarina R. Almeida

## Abstract

**Background:** Plasmacytoid dendritic cells (pDCs) play a key role in systemic sclerosis (SSc) pathophysiology. However, despite the recognised importance of metabolic reprogramming for pDC function, their metabolic profile in SSc remains to be elucidated. Thus, our study aimed to investigate the role of pDC metabolism in SSc.

**Methods:** Peripheral blood mononuclear cells (PBMCs) were isolated from the blood of healthy donors and SSc patients. SCENITH™, a single-cell flow cytometry-based method, was applied to infer the metabolic profile of circulating pDCs from patients with SSc. pDCs (CD304^+^ Lin^-^) at steady-state or stimulated with CpG A were analysed. Toll-like receptor (TLR)9 activation was confirmed by ribosomal protein S6 phosphorylation.

**Results:** Circulating pDCs from 10 healthy donors and 14 SSc patients were analysed. pDCs from anti-centromere antibody-positive (ACA^+^) patients displayed higher mitochondrial dependence and lower glycolytic capacity than those from anti-topoisomerase I antibody- positive (ATA^+^) patients. Furthermore, cells from both ACA^+^ patients and limited cutaneous SSc (lcSSc) patients showed a stronger response towards TLR9 activation than cells from ATA^+^, anti-RNA polymerase III antibody-positive (ARA^+^) or diffuse cutaneous SSc (dcSSc) patients.

**Conclusions:** Our results show that pDCs from ACA^+^ patients rely more on oxidative phosphorylation (OXPHOS) and are more responsive to external stimuli, suggesting that pDCs from ATA^+^ patients may be more activated or exhausted. These findings point for the possible contribution of pDC metabolism to SSc clinical course, unveiling new potential targets for therapeutic approaches in SSc.

## 1. Introduction

Systemic sclerosis (SSc) is a connective tissue disease characterised by immune dysregulation, vasculopathy and fibrosis of different organs [1]. SSc can be classified in two main subsets, based on the extent of skin involvement. In limited cutaneous SSc (lcSSc), fibrosis primarily affects the skin distal to the elbows or knees, whereas in diffuse cutaneous SSc (dcSSc), the involvement extends to the trunk [2,3]. Additionally, most SSc patients exhibit at least one specific serum autoantibody, namely anti-centromere, anti-topoisomerase I and anti-RNA polymerase III antibodies (ACA, ATA, ARA respectively), which are associated with disease presentation and progression [2,4]. ACA is typically associated with lcSSc, while ATA and ARA are more commonly found in dcSSc cases. Moreover, ATA^+^ patients are more prone to developing lung fibrosis, leading to a less favourable prognosis, whereas ARA^+^ patients often present renal crisis [2].

Plasmacytoid dendritic cells (pDCs) are professional type I interferon (IFN) producers involved in anti-viral responses, but also highly relevant for autoimmunity onset and flairs, particularly in diseases correlated with high IFN signature, such as SSc [5]. It has been reported that pDC numbers are increased in the skin and lung tissues of patients with SSc and decreased in circulation compared with healthy donors [6,7]. Also, pDC depletion in mice models of SSc ameliorates fibrosis and inflammation, further underlining their participation in disease progression [6,8]. pDCs can promote B cell activation and autoantibody production and secrete high levels of type I IFN and chemokine (C-X-C motif) ligand 4 (CXCL4), contributing to inflammation and fibrosis [7,9–11]. This situation is correlated with a dysregulation of the unfolded protein response (UPR) pathway inositol-requiring transmembrane kinase/endoribonuclease 1-alpha (IRE1α), that potentiates the tricarboxylic acid (TCA) cycle and was shown to contribute to the type I IFN signature observed in SSc patients [12].

Several immune cells, such as macrophages, dendritic cells (DCs), B cells and T cells, rely on metabolic reprogramming for differentiation and function [13]. Metabolic reprogramming of pDCs depends on the species and stimuli. Toll-like receptors (TLR)7/8-activated human pDCs were shown to upregulate oxidative phosphorylation (OXPHOS) dependently on glutaminolysis [14]. On the other hand, human pDCs stimulated with a TLR7 agonist, such as influenza A virus (IAV) or rhinovirus (RV), display increased glycolysis while OXPHOS was not impacted [15]. Other studies using mouse pDCs demonstrated that type I IFN derived from TLR9 activation contributes to fatty acid oxidation (FAO) and OXPHOS [16]. Most recently, it was proposed that chronic activation of pDCs in SSc could be due to dysregulation of metabolic pathways [12]. Nevertheless, the metabolic profile of these cells and its contribution to the disease’s progression is unknown. Thus, this work aimed to elucidate the metabolic profile of pDCs in SSc and investigate its possible contribution to disease course and prognosis. For that, we used SCENITH™, a single-cell technique that allows for the analysis of the metabolic profile of rare cell populations by flow cytometry, by assessing translation inhibition [17].

## 2. Methods

### 2.1. Patient inclusion and clinical data

Adult patients with SSc fulfilling the 2013 American College of Rheumatology/ European League Against Rheumatism classification criteria were included [4]. Patients with overlapping syndromes were not considered. All participants signed an informed consent form before inclusion and clinical data collected were anonymised. This study was approved by the Ethics Committee of Centro Hospitalar do Baixo Vouga (now ULS-RA) (Reference 44-069-2021).

Demographic, clinical and laboratory data were extracted from patients’ medical records. Demographic information included gender, birth date, tobacco and alcohol use. Disease onset was defined as the time of the first occurrence of a non-Raynaud’s symptom of SSc. Patients were categorised as lcSSc or dcSSc using LeRoy’s criteria [18]. Clinical data included SSc-related symptoms and signs (Raynaud’s phenomenon, digital ulcers, telangiectasia, puffy hands, sclerodactyly, calcinosis), modified Rodnan skin score (mRSS), and organ involvement. We used consensus definitions to characterise each major organ involvement of SSc, which were established in earlier studies [19]. We also included the nailfold videocapillaroscopy (NVC) pattern, the presence of comorbidities (arterial hypertension, diabetes mellitus, dyslipidaemia, and depression) and the ongoing treatments (glucocorticoids, immunomodulators, and vasodilators). Antinuclear autoantibodies (ANAs) were detected with an indirect immunofluorescence test (IIFT) on human epithelial cells (Hep-2) and anti– extractable nuclear antigen (ENA) analysis using the commercially available line immunoblot assay [EUROLINE Systemic Sclerosis (Nucleoli) Profile (IgG); Euroimmun, Lübeck, GER].

### 2.2. Cell isolation

Whole blood (20 mL) was collected in S-Monovette® Lithium Heparin (Sarstedt, Nümbrecht, GER) tubes, maintained at room temperature (RT) and processed within 1 hour of venepuncture. Following blood centrifugation at 400 xg at RT for 10 minutes and plasma recovery, peripheral blood mononucleated cells (PBMCs) were isolated by density gradient centrifugation. Briefly, blood was diluted twice in PBS (Gibco, Thermo Fisher Scientific, Waltham, MA, USA), overlaid onto Ficoll-Paque Plus (Cytiva, Marlborough, MA, USA) and cells were separated by centrifugation at 400 xg for 10 minutes at 4 °C in a swinging bucket rotor without brake. The mononuclear cell layer was transferred into a new tube, washed twice with PBS and finally with RPMI 1640 medium with 2 mM L-glutamine (Gibco).

### 2.3. Cell activation, SCENITH™ and pDC staining

To evaluate the metabolic profile of pDCs, we employed the flow cytometry based SCENITH™ method, which relies on the impact of metabolic pathway inhibition on protein synthesis levels to infer ATP and GTP availability [17]. SCENITH™ protocol was adapted from its previous description [17]. A scheme with the key steps of the protocol is presented in **Figure 1**. Initially, 1.5x10^6^ PBMCs were seeded in 170 µL RPMI 1640 with 2 mM L-glutamine (Gibco) supplemented with 10% heat-inactivated foetal bovine serum (FBS; Sigma-Aldrich, St. Louis, MO, USA) and 1% Penicillin-Streptomycin (Gibco) in a round-shaped bottom 96-well plate in the absence or presence of CpG A (3 µM; InvivoGen, San Diego, CA, USA) for 3 hours. After activation, cells were treated for 40 minutes with vehicle control (C; DMSO), 2-deoxy-glucose (2-DG; 100 mM), oligomycin (O; 1 µM), a combination of 2-DG and O (DGO), and puromycin (10 µg/mL). As a negative control and 15 minutes before puromycin treatment, harringtonine (H) was added (2 µg/mL). After two washing steps with ice-cold FACS buffer (2% FBS, 2 µM EDTA, PBS) and before surface staining, cells were incubated for 15 minutes at 4 °C in the dark with a combination of Human TruStain FcX Fc Receptor Blocking Solution (BioLegend, San Diego, CA, USA) and Live/Dead™ (Invitrogen, Thermo Fisher Scientific). Surface staining with primary conjugated antibodies was performed for 25 minutes at 4 °C in the dark. After washing with FACS buffer, cells were fixed and permeabilised using Foxp3 Transcription Factor Staining Buffer Set (Invitrogen), according to the manufacturer’s instructions. Cells were incubated for 10 minutes at RT with intracellular blocking solution [Foxp3 Permeabilization buffer (Invitrogen) with 20% FBS], and puromycin intracellular staining was performed for 1 hour at 4 °C. After two washing steps with permeabilization buffer, cells were resuspended in FACS buffer.

**Figure 1.**
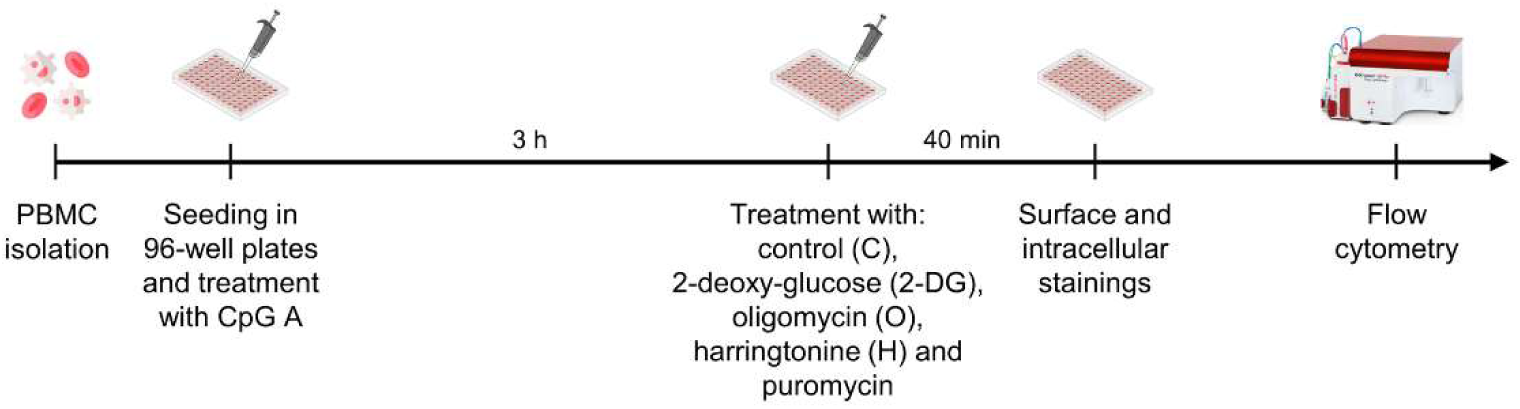
Scheme representing the key steps of the methods used. PBMCs isolated from the blood of HCs and SSc patients were seeded in 96-well plates and treated with CpG A for 3 hours, before 40 minute incubation with the SCENITH™ drugs – control (C), 2-deoxy-glucose (2-DG), oligomycin (O), harringtonine (H) and puromycin. Finally, stainings were performed and cells were analysed by flow cytometry.

The kit with SCENITH™ reagents [inhibitors, puromycin and anti-puromycin antibody) was from www.scenith.com. Information regarding the used antibodies is shown in **Supplementary Table S1**.

Data were acquired using BD Accuri C6 (BD Biosciences, San Jose, CA, USA). At least 300 CD304^+^ Lin^-^ events were acquired per condition. Data were analysed using FlowJo™ v10.8.1 Software (BD Life Sciences, Franklin Lakes, NJ, USA) and were compensated using single stains with UltraComp eBeads™ Compensation Beads (ref. 01-2222-41, Invitrogen). The gating strategy is shown in **Supplementary Figure S1**.

Glucose dependence, mitochondrial dependence, glycolytic capacity and fatty acid oxidation (FAO)/ amino acid oxidation (AAO) capacity were calculated as detailed below, using the median fluorescence intensities (MFI) of anti-puro-fluorochrome obtained upon treatment.

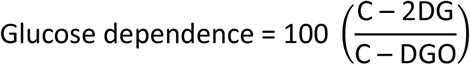

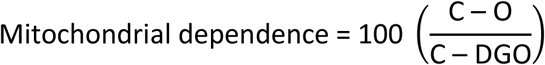

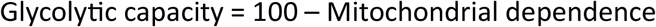

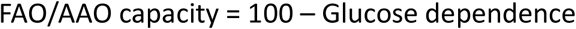

Glucose and mitochondrial dependences give the proportion of translation dependent on glucose oxidation or OXPHOS, respectively [17]. Glycolytic capacity represents the maximum translation sustainability under mitochondrial respiration inhibition, while FAO and AAO capacity represents cell’s ability to use fatty acids and amino acids as energy source when glucose oxidation is inhibited [17].

### 2.4. p-S6 staining

PBMCs were seeded, treated with CpG A and their surface stained as described for the SCENITH™ protocol. Cells were then stained for the intracellular phosphorylated form of ribosomal protein S6. Information about the used antibodies is in **Supplementary Table S1**.

### 2.5. Statistical analysis

Statistical analyses were performed using GraphPad Prism Software version 9.0.0 (GraphPad Software, Inc., La Jolla, CA, USA). Data are presented as median and interquartile range (IQR). The Shapiro-Wilk test was used to assess data normality. The most appropriate statistical test was then chosen according to each data set, as indicated in figure legends. *p < 0.05; **p < 0.01; ***p < 0.001; ****p < 0.0001.

## 3. Results

### 3.1. Study cohort characterisation

Fourteen SSc patients and ten healthy controls (HCs) were enrolled in this study (**Table 1**), with median ages of 56.5 (interquartile range, IQR 52.5-61.8) and 49 (IQR 35.8-53.0) years, respectively. Most were women (78.6% of SSc patients and 90% of HC). Median disease duration was 90 months (IQR 45-120). Eight patients (57.1%) presented lcSSc, and six (42.9%) presented dcSSc. Nine patients (69.2%) had gastrointestinal (GI) involvement (100% upper GI tract), and six (46.2%) presented interstitial lung disease (ILD). No patients with pulmonary arterial hypertension (PAH) or renal or cardiac involvement were included. Six patients (42.9%) were positive for ATA, four (28.6%) for ACA, and four (28.6%) for ARA. None of the patients was positive for more than one SSc-specific autoantibody.

**Table 1.** Participant demographics, clinical characteristics, and current therapies.

| Characteristic | SSc, n=14 | HCs, n=10 |
| --- | --- | --- |
| Gender (%) |  |  |
| Male | 3 (21.4) | 1 (10) |
| Female | 11 (78.6) | 9 (90) |
| Age (years) | 56.5 (52.5-61.8) | 49 (35.8-53.0) |
| Limited cutaneous SSc (%) | 8 (57.1) | - |
| Diffuse cutaneous SSc (%) | 6 (42.9) | - |
| Disease duration (months) | 90 (45-120) | - |
| mRSS | 11 (8-15) | - |
| Presence of autoantibodies |  |  |
| ATA (%) | 6 (42.9) | - |
| ACA (%) | 4 (28.6) | - |
| ARA (%) | 4 (28.6) | - |
| Organ complications |  |  |
| GI involvement (%) | 9 (69.2) | - |
| ILD (%) | 6 (46.2) | - |
| PAH (%) | 0 | - |
| Cardiac involvement | 0 | - |

| Current therapies |  |  |
| --- | --- | --- |
| Prednisolone (%) | 4 (30.8) | - |
| Methotrexate (%) | 4 (30.8) | - |
| Mycophenolate Mofetil (%) | 5 (38.5) | - |
| Nintedanib (%) | 2 (15.4) | - |
| Calcium antagonist (%) | (100) | - |

Median and IQR are reported unless otherwise stated. HC: healthy control; mRSS: modified Rodnan skin score; ATA: anti-topoisomerase I antibody; ACA: anti-centromere antibody; ARA: anti-RNA polymerase III antibody; GI: gastrointestinal; ILD: interstitial lung disease; PAH: pulmonary arterial hypertension.

### 3.2. The percentage of pDCs in circulation varies with disease duration and ANA titre

Treatments were performed on the whole PBMC population, and pDCs were identified by flow cytometry as positive for the pDC marker CD304 (also known as BDCA4) and negative for CD3, CD14, CD16, CD19, CD20, CD34 and CD56 (Lin^-^) (**Figure 2A**, gating strategy presented in **Supplementary Figure S1A**). It was confirmed that CD304^+^ cells are also positive for the pDC markers CD123 and HLA-DR, while events positive for these pDC markers are also Lin^-^ (**Figure S1B**, **C**).

**Figure 2.**
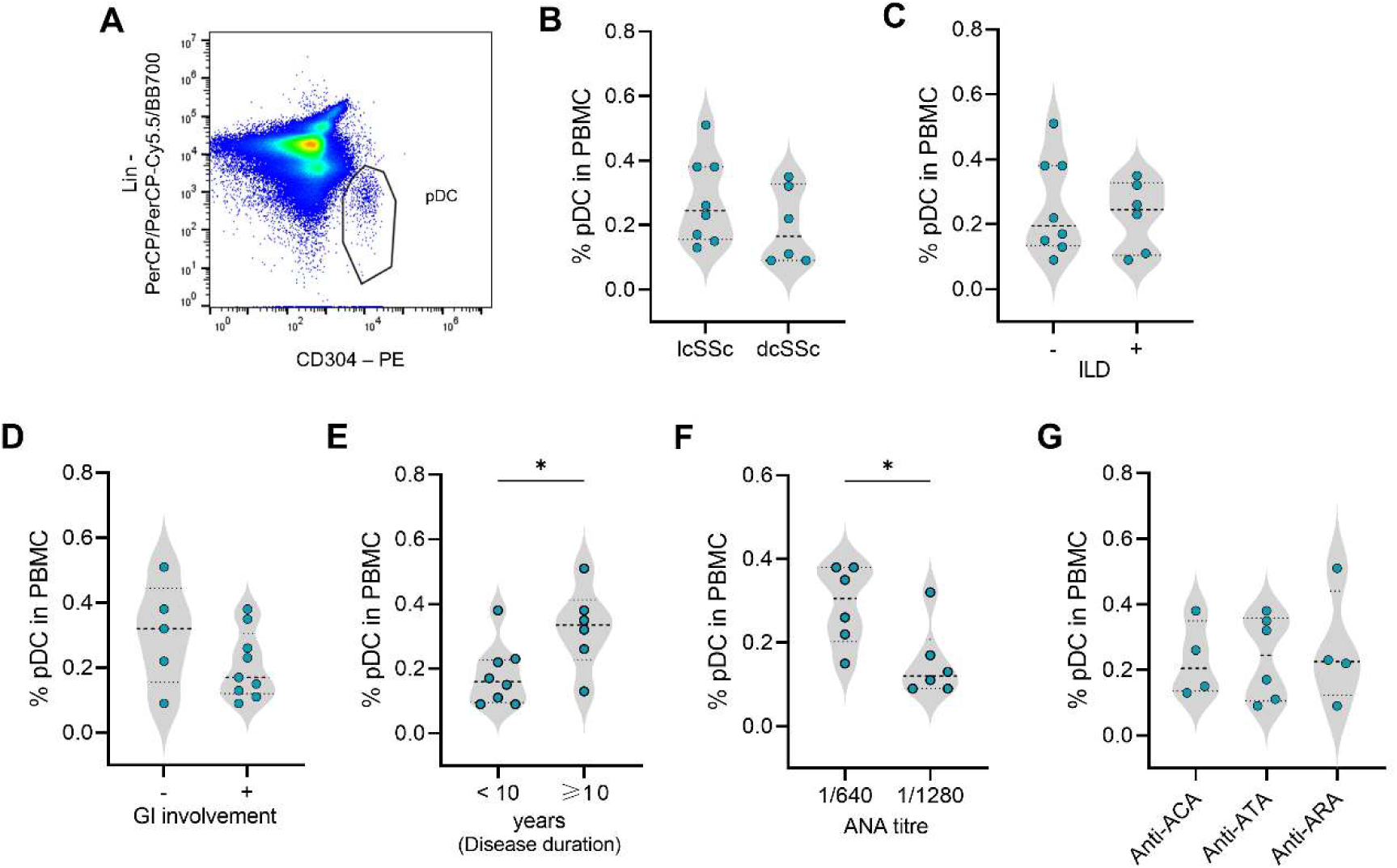
The percentage of circulating pDCs in SSc patients changes with progression of the disease. **(A)** Identification of CD304^+^ Lin^-^ pDCs by flow cytometry. **(B-G)** The percentage of pDCs was obtained by flow cytometry analysis and comparisons were performed between patient groups. Data are expressed as median and IQR and each dot represents data from one donor. Unpaired *t* test or one-way ANOVA followed by Tukey’s multiple comparisons test. *p< 0.05.

No statistically significant differences were found in the percentage of pDCs in circulation from patients with lcSSc or dcSSc (**Figure 2B**) and with or without ILD or GI involvement (**Figure 2C**, **D**). Nevertheless, patients diagnosed for at least ten years showed higher levels of pDCs in circulation (**Figure 2E**). Also, patients with higher ANA titre (1/1280) exhibited a reduced percentage of pDCs among PBMCs (**Figure 2F**), regardless of the subtype of autoantibody present (**Figure 2G**).

### 3.3. pDCs present a similar metabolic profile in SSc and HCs

Inhibition of glycolysis and OXPHOS by 2-deoxy-glucose (2-DG) and oligomycin (O), respectively, led to a reduction in translation to levels similar to harringtonine (H), a protein synthesis inhibitor (**Figure 3A**, **B**). This shows that pDCs are susceptible to metabolic modulation and rely on both glycolysis and OXPHOS for protein synthesis, measured by puromycilation detection and used as a read-out for ATP and GTP availability. Under basal conditions, pDCs from HCs and SSc patients showed similar translation and S6 phosphorylation levels (**Figure 3C**, **D**). As expected, TLR9 activation by CpG A led to increased levels of phosphorylated S6 in both HCs and SSc patients, even though statistical significance is only observed for the latter (**Figure 3D**). However, treatment with CpG A did not affect protein synthesis on pDCs, nor their response to metabolic modulation (**Figure 3A-C**). Moreover, the metabolic profile of pDCs was not affected by TLR9 activation and seemed comparable under both SSc and healthy conditions (**Figure 3E, F**). Given the lack of differences upon TLR9 stimulation, the follow-up analysis on the metabolic profile of pDCs was focused on cells in basal conditions.

**Figure 3.**
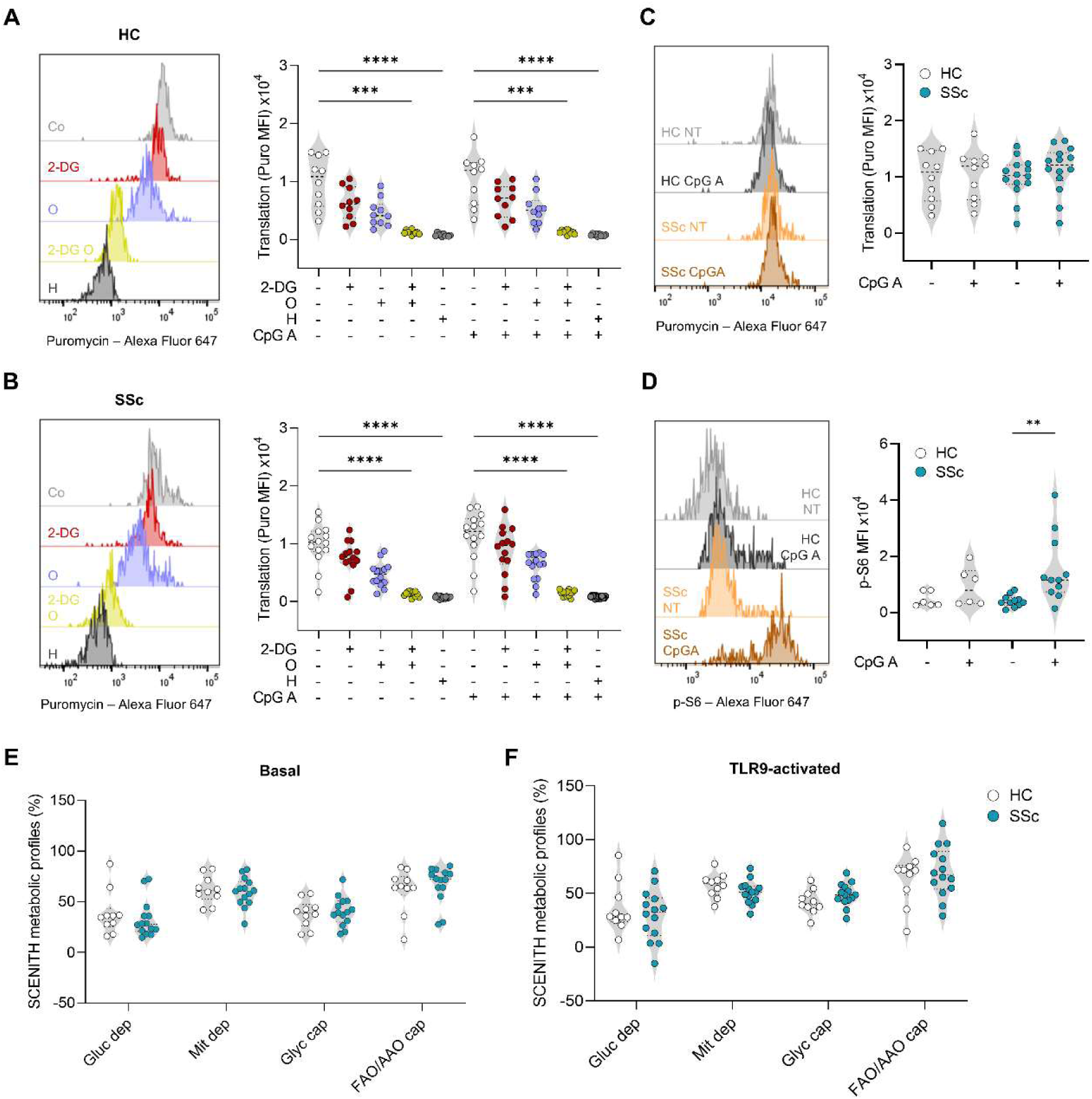
pDCs from HCs and SSc patients have no major differences in their metabolic profile. **(A, B)** Translation levels in pDCs after treatment with 2-deoxy-glucose (2-DG), oligomycin (O), harringtonine (H) and CpG A. Representative histograms for cells not TLR9-activated. **(C)** Translation and **(D)** S6 phosphorylation (MFI) levels. **(E, F)** SCENITH™ metabolic profiles. Data expressed as median and IQR; dots represent independent donors. Kruskal-Wallis test followed by Dunnet’s multiple comparisons test (A, B), one-way ANOVA followed by Sidak’s multiple comparisons test (C, D) and two-way ANOVA followed by Sidak’s multiple comparisons test (E, F) were used. ***p<0.001; ****p<0.0001.

### 3.4. pDCs from ACA and ATA-positive patients have different metabolic profiles

During our analysis we noted potential differences among groups of SSc patients and decided to segregate patients according to their clinical characteristics. pDCs from lcSSc patients display higher upregulation of S6 phosphorylation and protein synthesis after treatment with CpG A, in comparison with cells from dcSSc patients, suggesting higher sensitivity to TLR9 stimulation (**Figure 4A**, **B**). There was also a significant difference in the basal levels of translation associated with GI involvement but not with ILD, disease duration or ANA titre (Information about the used antibodies is in **Supplementary Figure S2**). However, at basal conditions, no differences were found in the metabolic profile of cells isolated from lcSSc or dcSSc (**Figure 4C**), and no correlation with either ANA titre, disease duration, GI involvement or ILD was found (**Supplementary Figure S3**).

**Figure 4.**
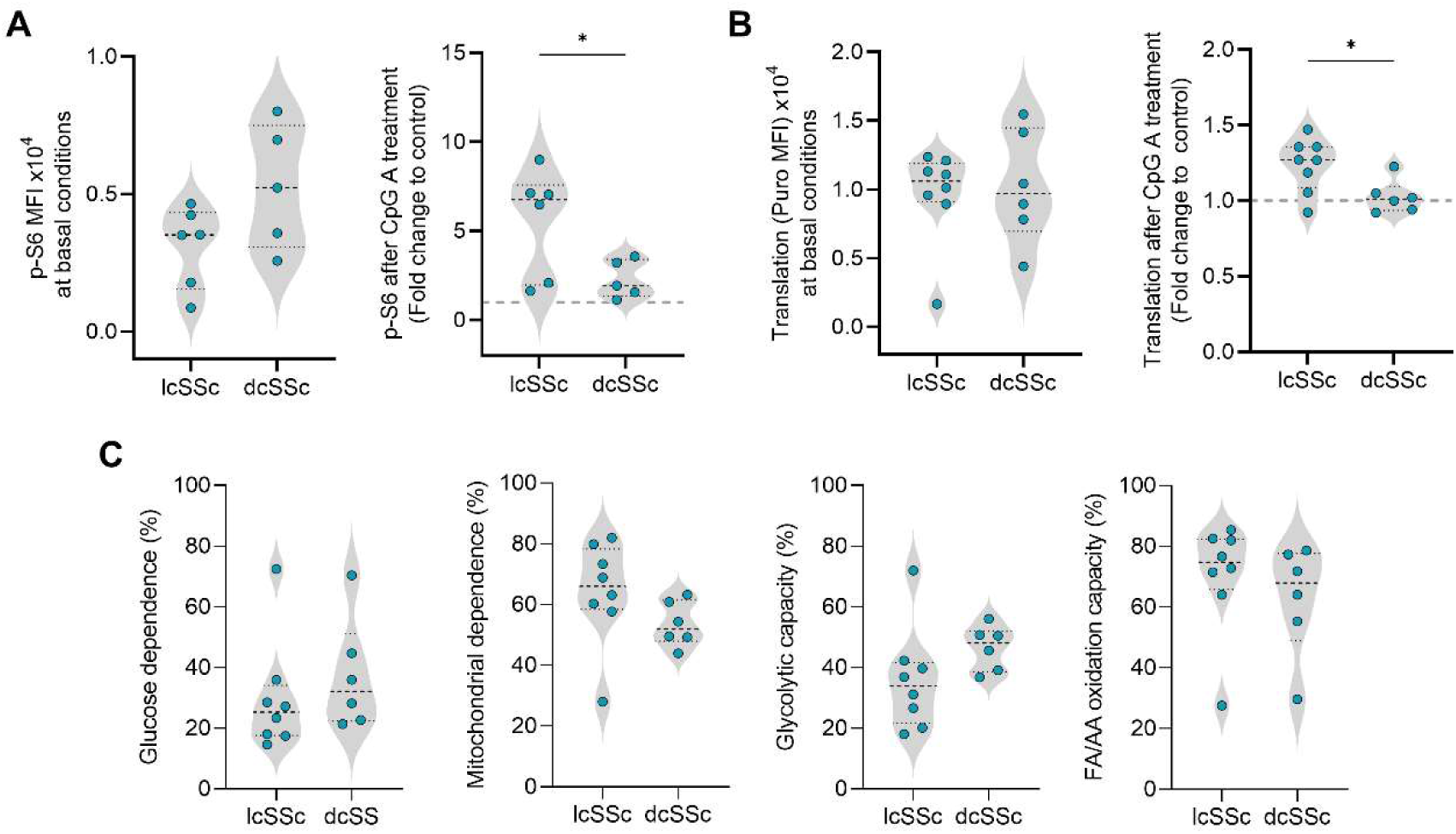
pDCs from lcSSc patients are more reactive to CpG A than cells from dsSSc. **(A)** p-S6 levels and **(B)** translation levels at basal and TLR9-activated conditions were assessed in pDC from SSc patients by flow cytometry. **(C)** Metabolic profiles. Dash line corresponds to 1. Data are expressed as median and IQR and each dot represents data from one donor. Statistical significance was tested with unpaired *t* test or Mann-Witney test. *p<0.05.

However, pDCs from ACA^+^ patients exhibited higher mitochondrial dependence and lower glycolytic capacity compared to cells from ATA^+^ patients (**Figure 5A**). Moreover, pDCs from ACA^+^ patients showed an upregulation of translation levels after TLR9 stimulation, compared to ATA^+^ and ARA^+^ patients (**Figure 5B**). There were no statistically significant differences on levels of p-S6, but there was a tendency for pDCs from ARA^+^ patients to display higher p-S6 levels at basal conditions (**Figure 5C**).

**Figure 5.**
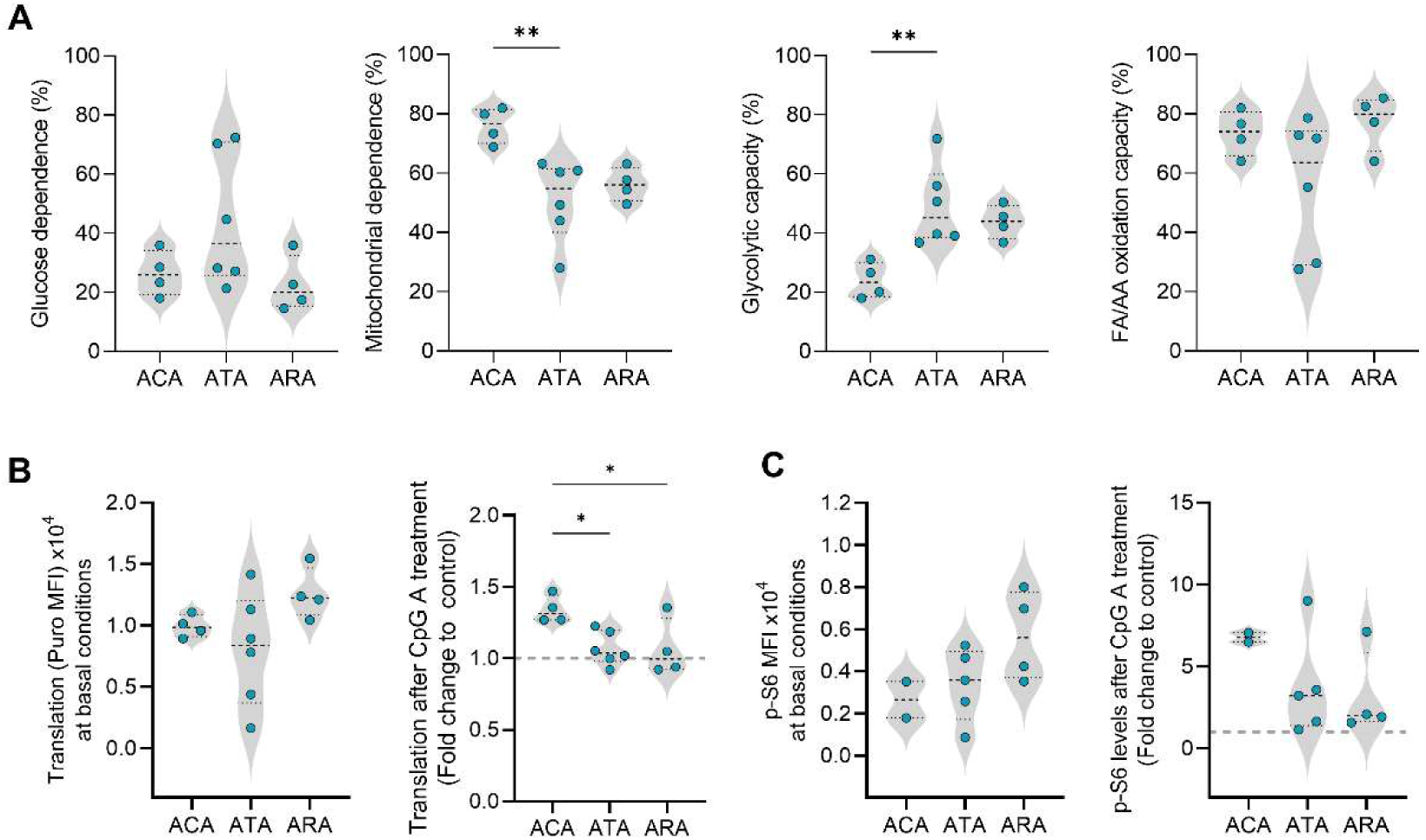
pDCs from ACA^+^ patients are more dependent on mitochondrial metabolism and have lower glycolytic capacity than cells from ATA^+^ patients. **(A)** SCENITH™ metabolic profiles, **(B)** translation levels, and **(C)** p-S6 levels were determined by flow cytometry for pDCs from patients with different patterns of SSc autoantibodies. Data are expressed as median and IQR and each dot represents data from one donor. Kruskal-Wallis test followed by Dunnet’s multiple comparisons test was used for (A), one-way ANOVA followed by Tukey’s multiple comparisons test for (B) and one-way ANOVA followed by Sidak’s multiple comparisons test for (C). *p< 0.05, **p<0.01.

## 4. Discussion

The key role of pDCs in SSc pathphysiology has been highlighted by others [7,10–12], primarily using mouse models of scleroderma that revealed an improvement in fibrosis upon pDC depletion [6,8]. Furthermore, pDCs are hyperactivated in patients with SSc, contributing to the characteristic type I IFN signature [7,9,10]. As with other immune cells, pDCs heavily rely on metabolic reprogramming for their differentiation and function [20,21]. A reduced expression of phosphoglycerate dehydrogenase (PHGDH) in pDCs isolated from the blood of SSc patients, and the downregulation of interferon-stimulated genes (ISGs) expression upon TCA cycle inhibition were recently reported [12]. Nevertheless, the metabolic profile of pDCs in SSc has not been characterised. In this study, we used SCENITH™, a single-cell resolution method that employs flow cytometry analysis of protein synthesis to infer the metabolic profile of cells [17]. The reliance of cells on glucose oxidation and OXPHOS for the maintenance of protein synthesis levels is given by glucose and mitochondrial dependences, respectively. Simultaneously, the ability of cells to sustain mRNA translation under conditions of mitochondrial respiration or glucose oxidation inhibition is reflected in the glycolytic and FAO/AAO capacities, respectively. Due to the importance of cell activation in this context, we analysed both cells at basal conditions and pre-treated with CpG A, a TLR9 agonist. Cell activation was monitored by phosphorylation of S6, which occurs upon TLR9 activation [22]. Treatments were performed in the whole PBMC population, and the analysis focused on CD304^+^ Lin^-^ cells.

The identification of pDCs by flow cytometry as CD304^+^ Lin^-^ (CD3, CD14, CD16, CD19, CD20, CD34 and CD56) can be viewed as a possible limitation of this work. Even though we confirmed that these cells were also positive for other pDC markers (CD123 and HLA-DR), we cannot certainly exclude a low percentage contamination by other cells. Moreover, these findings are in circulating pDCs, which may not be the cell population with the highest influence on the disease course. Infiltrating pDCs likely have a more prominent role in SSc pathogenesis. However, as SSc is a systemic disease, it is also possible that the profile of circulating pDCs reflects what happens in the affected tissues. Finally, our analyses were performed with a low number of cells and patients, due to the rarity of this cell population and disease. However, despite the low number of patients included, the sex and age distribution of the patients included reflects the Portuguese SSc patients’ demographics [23].

Patients diagnosed for less than ten years at the time of blood collection displayed lower amounts of pDCs in circulation than those diagnosed for longer. Typically, pDCs circulate through the blood and are recruited to target tissues upon infection or inflammation [24]. SSc is characterised by an initial vascular injury and early inflammation phase that contribute to fibrosis development [25,26]. This difference in pDC levels may be due to the early recruitment of pDCs to the inflamed tissues, a phenomenon that could decrease as the disease progresses. Also, patients with higher levels of ANAs in circulation displayed less pDC abundance within PBMCs. ANA titres are suggested to be associated with disease progression, and in some cases, high titres are correlated with the disease outcome [27]. Higher ANA titres may indicate a more robust autoimmune response and inflammation, possibly justifying these results [28,29]. However, this is a debated theory.

The activation state of pDCs was evaluated by analysing p-S6 levels and monitoring protein synthesis, which is reported to increase during immune cell activation to sustain cellular function [30–32]. Globally, pDCs from HCs and SSc patients seemed to have the same basal activation state since no differences in p-S6 and translation levels were found. However, within the patient groups, differences were identified among patients with GI involvement, displaying lower translation levels compared to patients without this involvement. The exact changes that occur in the GI tract are not fully characterised in SSc [33], making it difficult to find a justification for this result. However, it may be speculated that there might be sequestration of cells with higher translation rates in the affected tissues and/or disparities in the microbiome. In response to TLR9 activation, pDCs from ACA^+^ patients upregulated their translation levels, contrary to what happens with ATA^+^ or ARA^+^ patients. ATA and ARA are more common in patients with dcSSc, while ACA is often associated with lcSSc. A higher increase in translation upon TLR9 activation was also observed in lcSSc, compared to dcSSc. Although not statistically significant, there was a tendency for pDCs from dcSSc and patients positive for ATA or ARA to present increased p-S6 at basal conditions. Therefore, this suggests that pDCs from dcSSc and ATA^+^ and ARA^+^ patients have basal activation, which might limit the effect of CpG A.

Even though the treatment with CpG A induced S6 phosphorylation, it did not upregulate translation, nor impact the metabolic profile of these cells. Other studies have observed an impact of TLR9 activation in the metabolism of pDCs [16,34], but longer exposure times to the agonist were used. As mentioned above, a more recent study exploring the metabolism of pDCs from SSc patients used a similar timepoint and analysed the expression of an enzyme involved in the biosynthesis of serine [12]. Therefore, differences from what is reported in the literature may be due to differences in the agonist and stimulation times used.

Interestingly, although pDCs from HCs and SSc patients displayed similar metabolic profiles, being dependent on both glycolysis and OXPHOS, variations were found among patients positive for the different ANAs subtypes. While there were no differences in the activation or metabolic state of pDCs from ATA^+^ and ARA^+^ patients, pDCs from ACA^+^ patients presented higher mitochondrial dependence and lower glycolytic capacity than those from ATA^+^ patients. This suggests that pDCs from ACA^+^ patients rely more on OXPHOS for energy production. On the other hand, cells from ATA^+^ patients tended to be more dependent on glycolysis. Usually, activated immune cells and highly proliferating cells shift their metabolism to glycolysis, while OXPHOS is preferred at the resolution phase [35]. Thus, the metabolic profile results are concordant with p-S6 readouts and suggest that pDCs from ATA^+^ patients are more activated, relying on glycolysis for energy production. These results are in accordance with the observations for pDCs from lcSSc (which are more commonly ACA^+^) compared to dcSSc patients. On the other hand, although both ATA and ARA are associated with dcSSc, the clinical course differs. While ATA^+^ patients are more prone to develop ILD, ARA is associated with scleroderma renal crisis and the co-occurrence of cancer [2,27].

The interaction between autoantibodies and soluble antigens can form immune complexes (ICs), promoting cell activation. Particularly, ICs containing antibodies specific for SSc (SSc-ICs) were described to promote the activation of skin fibroblasts and endothelial cells from HCs, having a pro-inflammatory and pro-fibrotic effect [36,37]. Furthermore, the mixture of sera from SSc patients with necrotic/apoptotic material induces pDC activation, apparently by SSc- ICs [38]. Even though not every patient’s serum can form ICs capable of inducing an IFN response, this phenomenon was observed with samples from both lcSSc and dcSSc [38]. In these studies, the autoantibody composition analyses did not always include the ones present in this work, and the formation of interferogenic ICs in ATA^+^ sera is controversial [38,39]. Thus, it would be interesting to clarify if the differences in the metabolic profile of pDCs and their activation status observed with our data are related to the activation of pDCs by ICs.

The presence of these different autoantibodies is linked with disease progression, and ATA^+^ patients tend to have a less favourable prognosis compared to those positive for ACA, partly due to the elevated risk of developing ILD, a common cause of death. Crucially, we found that pDCs from ACA^+^ patients had reduced glycolytic capacity, associated with a stronger response to *in vitro* TLR9 activation. Together, these results suggest that pDCs more dependent on mitochondrial respiration and less on glycolysis are associated with reduced basal activation and a better prognosis. Thus, we believe that our findings pave the way for understanding the potential involvement of pDC metabolism in the severity and clinical progression of SSc. Future testing of the impact of blocking glycolysis while promoting mitochondrial respiration on pDC may contribute to defining new possible therapeutic strategies for SSc.

## Acknowledgements

We thank all the volunteers, especially the patients, and the health professionals (Anabela Barcelos, Maria do Céu Morais, Graça Costa, António José Oliveira) involved in this study.

## CRediT authorship contribution statement

**Beatriz H. Ferreira:** Conceptualization, Data curation, Formal analysis, Investigation, Methodology, Visualization, Writing – original draft, Writing – review and editing. **Carolina Mazeda:** Conceptualization, Data curation, Investigation, Methodology, Resources, Writing – review and editing. **Eduardo Dourado:** Data curation, Methodology, Writing – review and editing. **João L. Simões:** Investigation, Methodology, Resources, Writing – review and editing. **Ana R. Prata:** Data curation, Methodology, Writing – review and editing. **Rafael J. Argüello:** Methodology, Resources, Writing – review and editing. **Iola F. Duarte:** Funding acquisition, Methodology, Supervision, Writing – review and editing. **Philippe Pierre:** Funding acquisition, Methodology, Supervision, Writing – review and editing. **Catarina R. Almeida:** Conceptualization, Funding acquisition, Investigation, Methodology, Project administration, Supervision, Writing – original draft, Writing – review and editing.

## Funding

This work was supported by the World Scleroderma Foundation and Edith Busch Stiftung (the EBF and WSF Research Grant Programme 2022-2023). Work developed within the scope of iBiMED – Institute of Biomedicine(UIDB/04501/2020 and UIDP/04501/2020) and CICECO – Aveiro Institute of Materials UIDB/50011/2020 (DOI 10.54499/UIDB/50011/2020), UIDP/50011/2020 (DOI 10.54499/UIDP/50011/2020) & LA/P/0006/2020 (DOI 10.54499/LA/P/0006/2020) and the project with the reference 2022.03217.PTDC (DOI 10.54499/2022.03217.PTDC), financially supported by national funds (OE), through Fundação para a Ciência e a Tecnologia (FCT)/MCTES. FCT is acknowledged for the individual grant to B.H.F. (SFRH/BD/144706/2019; DOI 10.54499/SFRH/BD/144706/2019) and the research contract under the Scientific Employment Stimulus to I.F.D. (CEECIND/02387/2018).

## Conflict of interest statement

No competing interests.

## Data availability statement

Data are available upon reasonable request.

## Supplementary material

**Supplementary Figure S1.**
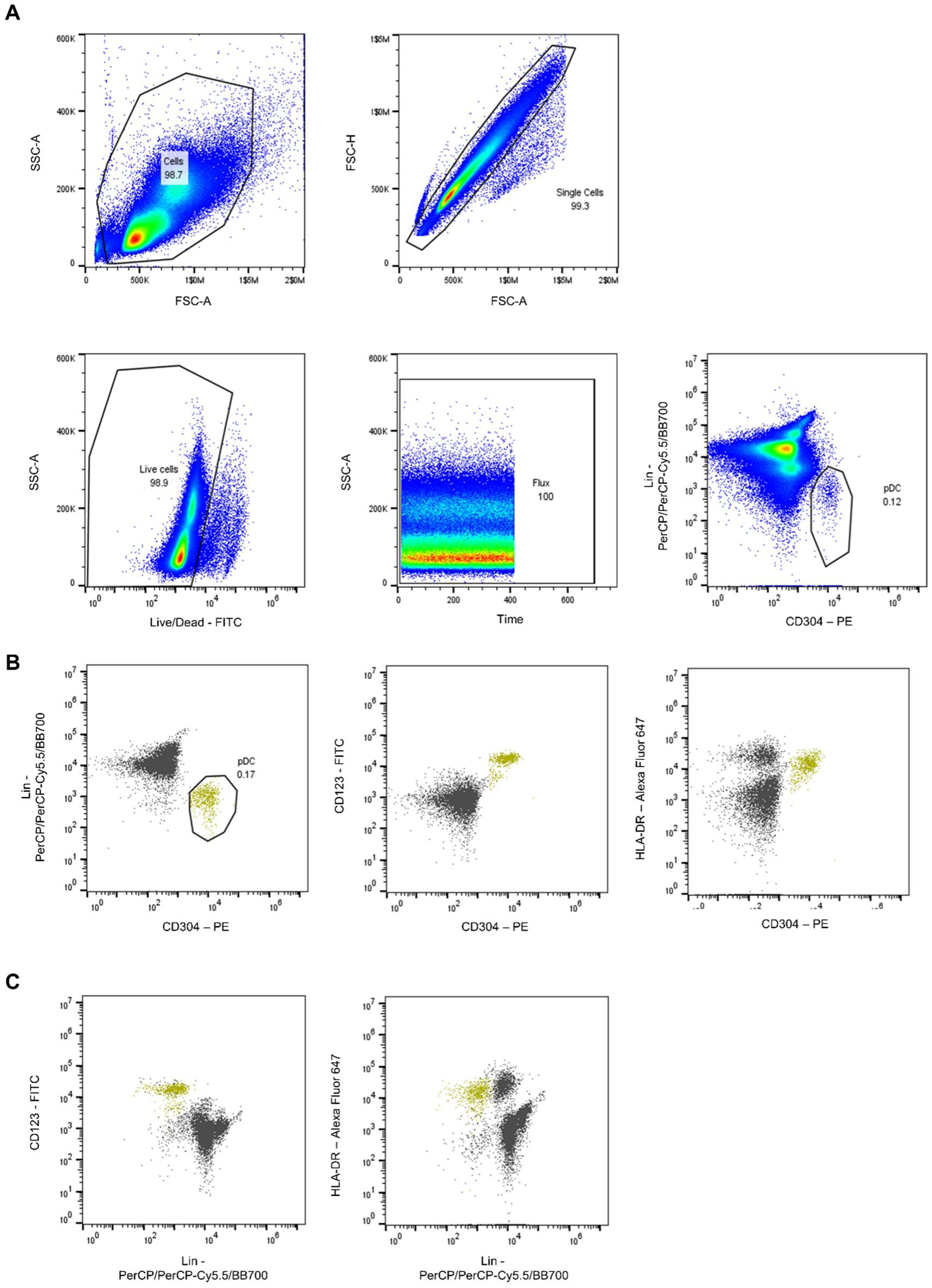
Gating strategy used. **(A)** Gating strategy for identification of pDCs as CD304^+^ Lin^-^ (CD3^-^ CD14^-^ CD16^-^ CD19^-^ CD20^-^ CD34^-^ CD56^-^). Only live cells acquired during stable flux were included in our analyses. **(B, C)** Confirmation that gating on CD304^+^ Lin^-^ (shown in yellow) allows identification of pDCs (which are also CD123^+^ and HLA-DR^+^).

**Supplementary Figure S2.**
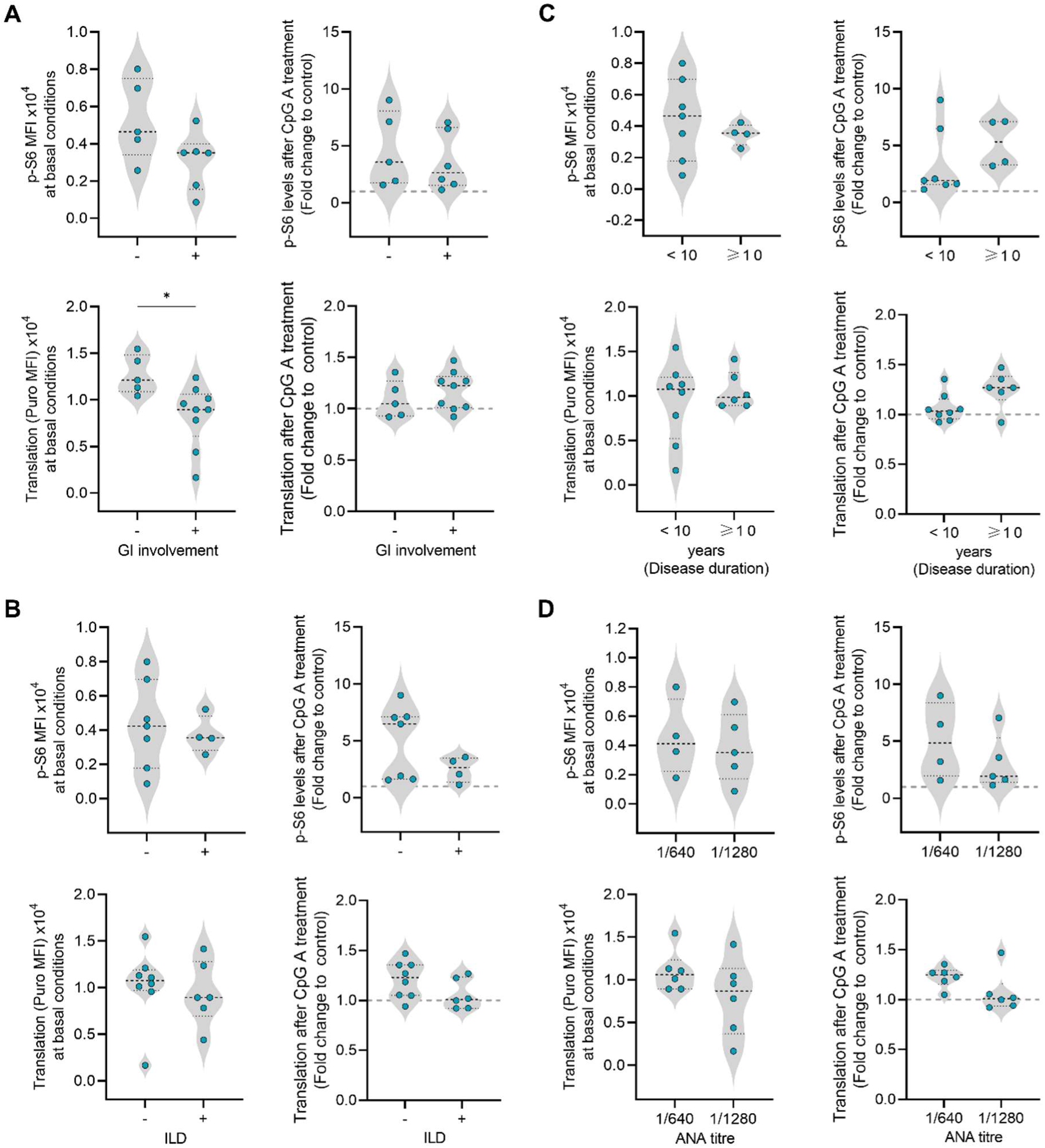
S6 phosphorylation and translation levels of pDCs from SSc patients according to their clinical data. **(A)** Gastrointestinal (GI) involvement, **(B)** interstitial lung disease (ILD), **(C)** disease duration or **(D)** ANA titre. Grey line corresponds to 1. Data are expressed as median and IQR and each dot represents data from one donor. Statistical significance was tested with unpaired *t* test or Mann-Witney test. *p<0.05.

**Supplementary Figure S3.**
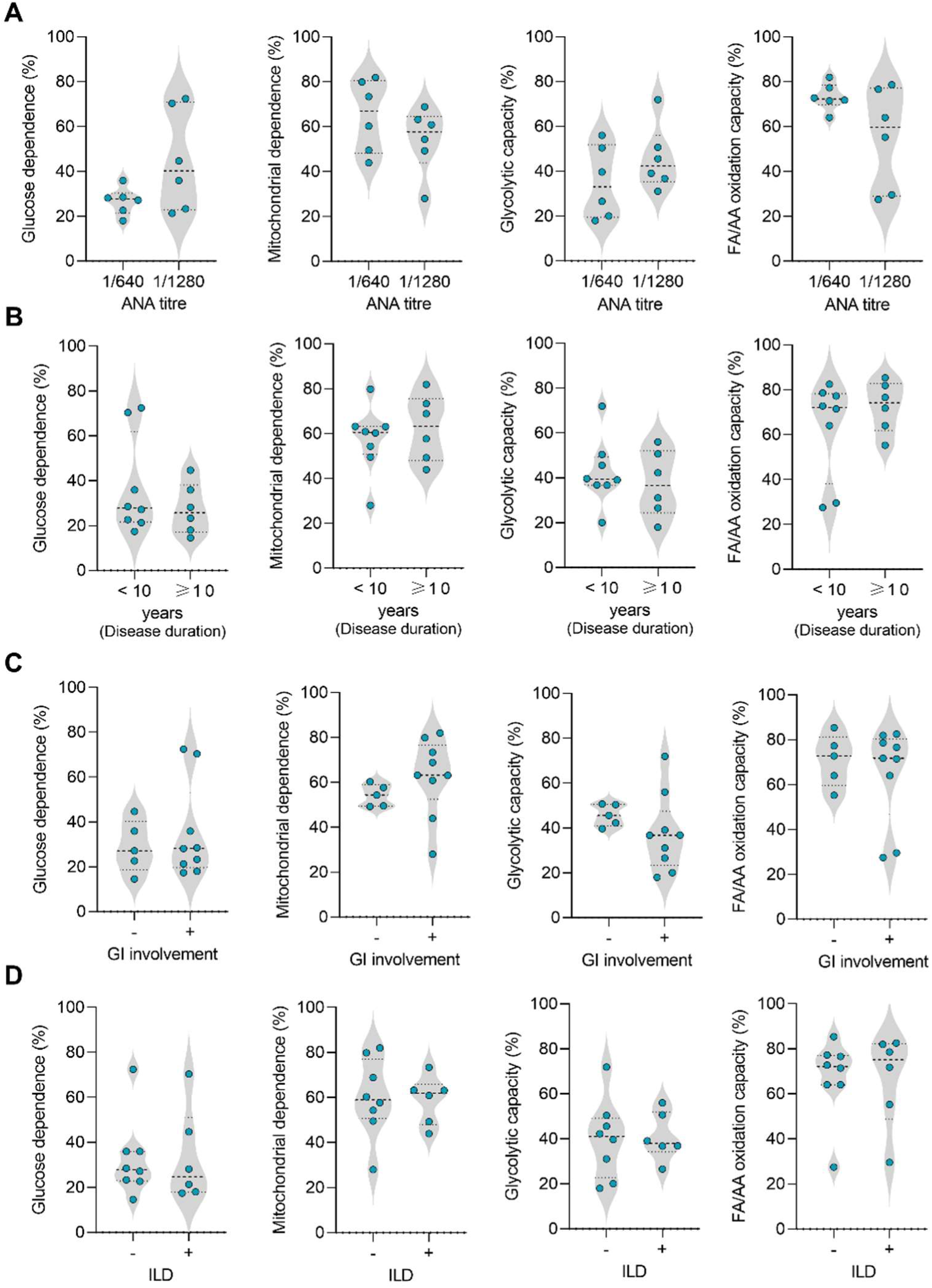
Metabolic profiles of pDC from SSc patients accordingly to their clinical data. **(A)** ANA titer, **(B)** disease duration, **(C)** gastrointestinal (GI) involvement, or **(D)** interstitial lung disease (ILD). Data are expressed as median and IQR and each dot represents data from one donor. Statistical significance was tested with unpaired *t* test or Mann-Witney test.

**Supplementary Table S1.** List of antibodies used in this study.

| **Antibody** | **Concentration** | **Reference** | **Source** |
| --- | --- | --- | --- |
| Anti-human CD304 PE (Clone: 12C2) | 2.5 µL/condition | 354504 | BioLegend |
| Anti-human CD123 FITC (Clone: 6H6) | 1.25 µL/condition | 306013 | BioLegend |
| Anti-human HLA-DR Alexa Fluor 647 (Clone: L243) | 2.5 µL/condition | 307621 | BioLegend |
| Anti-human CD3 PerCP (Clone: SK7) | 2.5 µL/condition | 344814 | BioLegend |
| Anti-human CD14 PerCP (Clone: M5E2) | 2.5 µL/condition | 301848 | BioLegend |
| Anti-human CD16 PerCP (Clone 3G8) | 1.25 µL/condition | 302030 | BioLegend |
| Anti-human CD19 PerCP (Clone: SJ25C) | 1.25 µL/condition | 363014 | BioLegend |
| Anti-human CD20 PerCP (Clone: 2H7) | 2.5 µL/condition | 302324 | BioLegend |
| Anti-human CD34 PerCP-Cy 5.5 (Clone: 561) | 5 µL/condition | 343612 | BioLegend |
| Anti-human CD56 BB700 (Clone: NCAM16.2) | 1.25 µL/condition | 566573 | BD Biosciences |
| Anti-puromycin Alexa Fluor 647 (Clone: R4743L-E8) | 1:100 | RRID:<br>AB_2827926 | [17] |
| Anti-phospho-S6 ribosomal protein (Ser235/236) | 1:50 | 2211 | Cell Signaling Technology |
| Anti-rabbit IgG Alexa Fluor 647 | 1:5000 | A-21245 | Invitrogen |

## Notes

### Competing Interest Statement

The authors have declared no competing interest.

